# Anatomical and functional examination of superior colliculus projections to the inferior olivary neurons in mice

**DOI:** 10.1101/2024.11.29.625963

**Authors:** Deviana David, Hugo Nusselder, Marylka Yoe Uusisaari

## Abstract

The Inferior olive (IO) is an important region for motor learning and movement coordination. Its climbing fibers projection to the Purkinje neurons in the cerebellar cortex is a sole source of the complex spikes, characterized by a strong depolarization in the Purkinje neuron’s dendrites. To generate spikes, the IO relies on inputs from various regions of the brain, including the superior colliculus (SC), a midbrain structure known for its role in orienting behaviors.

This study investigates SC projections to the IO using viral tracers, calcium imaging, and optogenetic stimulation. We reveal that, in addition to the known projections to the medial accessory olive (MAO), SC axons also project to the ventral principal olive (PO). Despite projecting to different parts of the IO, SC-MAO and SC-PO neurons are intermingled within the lateral part of the SC with similar gross morphology. We show that SC axons terminate on both dendritic shafts and spines of IO neurons, potentially influencing not only spiking probability, but also the network synchronization mediated by gap junctions coupling on the dendritic spines.

As a proof of principle, we recorded the in-vivo activity of neurons in ventral PO using calcium indicators and show that optogenetic activation of SC inputs can evoke spiking and enhance synchronization in IO neurons.

## 1 Introduction

The inferior olive (IO) is part of the olivocerebellar system (OCS), which is important for motor learning and coordinated movement. The OCS consists of the interconnected network between the IO, the cerebellar cortex, and the cerebellar nuclei (CN). The IO sends its projections, known as climbing fibers (CFs), to Purkinje neurons (PNs) in the cerebellar cortex. CF activity induces strong depolarization in the PN dendrites known as complex spikes (CSs), considered a key element of cerebellar function and motor learning (De Gruijl et al (2013); Streng et al (2018); Silva et al (2024); Medina and Lisberger (2008); Wagner et al (2021)). Collaterals of the CFs from IO also target the CN, whose neurons form the final stage of cerebellar computation.

Unlike the many types of spontaneous pacemaker cells in the OCS, the IO neurons’ spiking is driven by afferent inputs. The afferents of various regions of the brain converge in single cells (Ju et al (2019)), suggesting integrative processing within the IO. However, even though the combined effects of undefined excitatory inputs and GABAergic feedback from the cerebellum have been investigated (Loyola et al (2023)), our understanding of the specific termination patterns of identified afferent pathways remains low.

Among the various afferents, the IO receives afferents from the midbrain region called the superior colliculus (SC). The SC (known as the optic tectum in the non-mammalian organism) is an evolutionarily conserved region that plays an important role in behavioral orientation (Zhou et al (2023); Masullo et al (2019); Solié et al (2022); Basso and May (2017)). The layered structure of the SC is known to implement a spatial map of the external world, so that the anteroposterior (A-P) axis of the SC represents the front and rear visual field, while the mediolateral axis of the SC represents the upper and lower visual field (Cang et al (2018); Ito and Feldheim (2018)).The projections from the SC diverge into numerous motor and non-motor regions (Benavidez et al (2021)) that make use of the information about external world.

One of the less-investigated projections of the SC targets the IO. Studies across species, such as macaques, cats, rats, and mice, have demonstrated that SC projects to the medial accessory olive (MAO; Kyuhou and Matsuzaki (1991); Akaike (1985); T.Hess (1982); May et al (2023)). The fact that the neurons in MAO project to the paleocerebellum (cerebellar vermis; Sugihara and Shinoda (2004)) suggests that the tectal input could have had a significant value as one of the first inputs to the IO(Northcutt (2002). Thus, the SC-IO connection can be seen as an excellent starting point to examine multisensory integration in the IO.

A recent study (Benavidez et al (2021)) revealed a mediolateral segregation of SC axons in MAO. However, in order to consider the manner in which SC signals could be integrated with the ongoing activity within the IO network, the subcellular organization of the SC axons within the IO should be examined. Here, we further investigated the SC-IO pathway at both the mesoscopic and subcellular levels. First, using anterograde and retrograde viral labeling, we mapped the SC-IO projection across the IO and SC regions. Surprisingly, we found that in addition to the previously known targeting of the MAO, the principal olive (PO) region also receives axons from the IO. The PO-targeting SC neurons are likely distinct from the MAO-targeting ones, but intermingled in the same regions of the SC. Next, we examined the neurotransmitter identity of the SC-MAO and SC-PO axons, as well as mapped the location of axonal terminals along the dendrites of IO neurons. Intriguingly, we found that the SC axons target both dendritic shafts and spines, suggesting that activity in SC might not only drive spiking in IO neurons but also modulate levels of network synchronization that depend on electrotonic coupling via dendritic spines(Lefler et al (2014); Leznik and Llinás (2005); De Zeeuw et al (1998)). To examine this, we took advantage of the fact that the ventral fold of PO, targeted by SC axons, is accessible for GRIN-lens-based calcium imaging from the ventral side (Guo et al (2021)), and found evidence that the SC-IO pathway can evoke spikes as well as promote network-level synchronization. The work provides anatomical foundation for further studies of the SC-IO pathway in behaving animals.

## 2 Results

### 2.1 Dual projection of superior colliculus neurons into principal olive and medial accessory olive

In order to visualize the IO regions targeted by the superior colliculus (SC), we anterogradely labeled SC axons using a tdTomato-coupled viral tracer (Figure 1A-C). In addition to the well-established projection to the medial accessory olive (MAO), we discovered a smaller projection to the principal olivary neurons (PO). To explore the possibility that SC neurons projecting to MAO and PO originate from distinct SC regions, we employed two viral tracers tagged with separate fluorophores, delivered into medial and lateral regions of the SC. This method allowed us to distinctly mark populations of SC neurons and their axonal pathways to the inferior olive (Figure 1D). An example 10x confocal microscopy scan of the midbrain with labeled SC neurons in different medio-lateral (M-L) positions (out of 3) is shown in (Figure 1E). Figures 1 E1-E2 show the expression of tdTomato-labeled neurons in the medial and lateral parts of SC and the expression of eGFP in the lateral part of SC, with only a small fraction of the lateral SC colabeled (white region in Figure 1E1-E2). The example 20x scan of the medulla containing the IO in Figure 1 F1 shows SC axons labeled with tdTomato and GFP (Figure 1 F2) in contralateral subnucleus d of medial accessory olive (cdMAO). Here, tdTomato expression is seen closer to the midline, while eGFP expression are more towards the lateral part of cdMAO with a co-labeled area in the middle(Figure 1F3). Thus, medial SC neurons project to the medial part of the cdMAO while lateral SC projects to the lateral cdMAO. This SC-MAO projection aligns with the SC-IO projection pattern reported in previous findings (Benavidez et al (2021)). The ipsilateral projection of SC-MAO is weaker than the contralateral projection (Figure 1F).

**Fig. 1.**
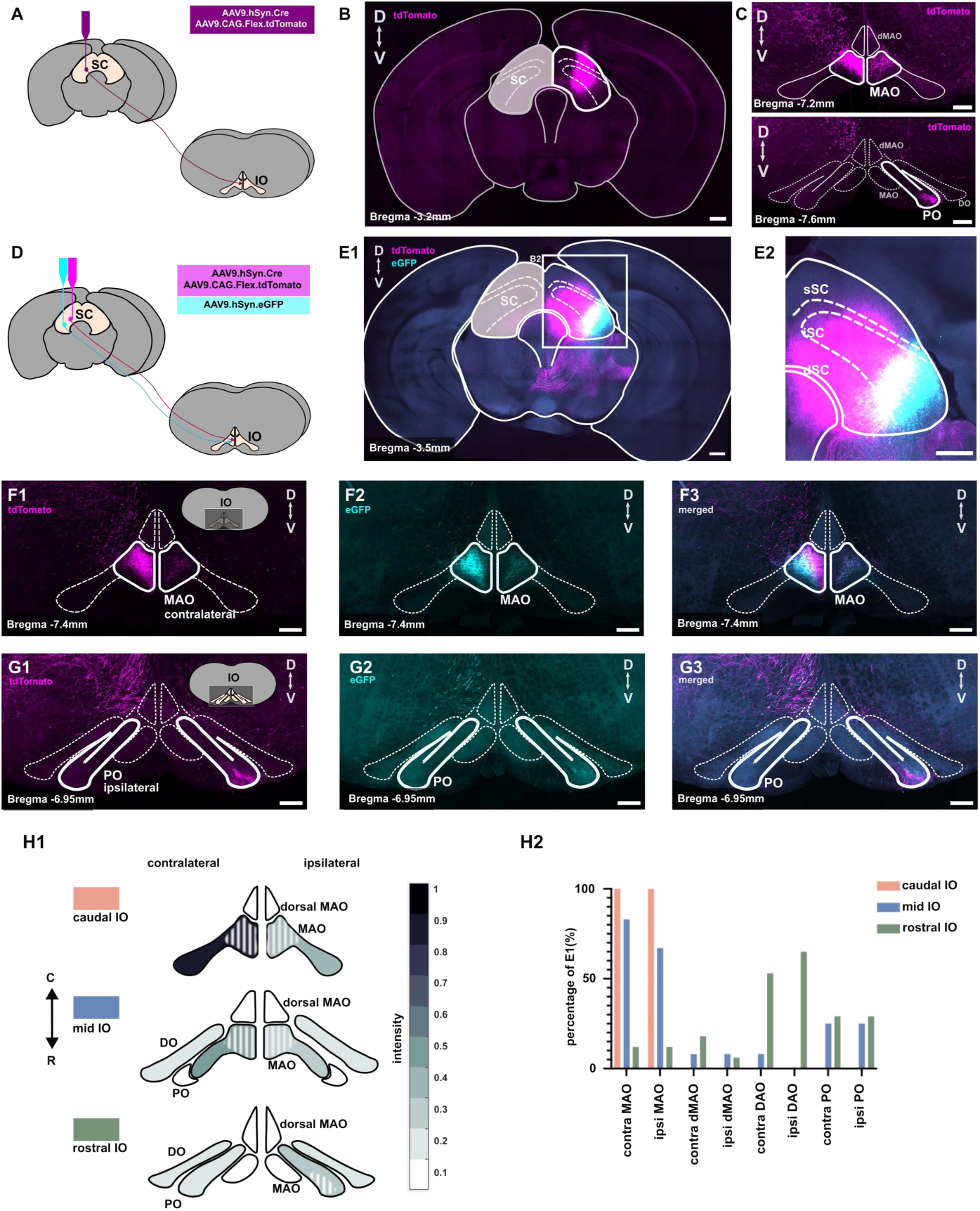
Superior colliculus (SC) projections to the medial accessory olive (MAO) and principal olive (PO). (**A**) Schematic of the injection of a mixture of AAV9.hSyn.Cre and AAV9.Flox.tdTomato to anterogradely label SC neurons, illustrated alongside a 10x confocal image of the midbrain and SC (**B**). Anterograde labeling reveals axons projecting not only to the MAO but also to the PO on the ipsilateral side (**C**). (**D**) Schematic showing the injection of a mixture of AAV9.hSyn.Cre, AAV9.Flox.tdTomato, and AAV9.hSyn.eGFP into different regions of the SC, resulting in labeled neurons within the medial and lateral SC, as depicted in (**E**). (**F1-3**) Images showing labeled SC axons in the contralateral and ipsilateral MAO, with another labeled axons found in the ipsilateral PO at the rostral part of the inferior olive (**G1-3**). (**H1**) Normalized density of labeled SC axons in diffent IO subnuclei along the rostro-caudal axis, with coordinates ranging from -6.9 mm to -7.9 mm from bregma. (**H2**) Probability distribution of finding labeled SC axons across different IO subnuclei, as shown in panel H1. Scale bar: : 200 *µ*m.

During careful examination of the SC-IO termination zones across all IO regions, we also found labeled axons in a small area corresponding to the ventral fold of the principal olive (PO). Figure 1G shows a 20x scan of the rostral medulla containing IO with labeled axons in the ipsilateral part of the PO (Figure 1G1-G3). While both tdTomato and eGFP were seen, the tdTomato labeling was much stronger,suggesting that SC-PO neurons are mainly located in the SC area that we targeted for tdTomato expression (Figure 1 E).

To summarize our findings in 5 animals, Figure 1H1 shows the normalized average intensity of fluorescence labeling in 48 IO slices across the rostro-caudal axis of the IO. Most SC-IO axons can be found in the caudomedial MAO, and in the rostral ipsilateral PO. The shaded area shows the location of the labeled axons within MAO and PO. The percentage of having labeled SC-IO axons as shown in Figure 1H1 in 5 animals is shown in the panel (Figure 1H2).

### 2.2 Superior colliculus neurons projecting to the medial and principal olive are intermingled

To provide additional insight into the organization of the SC-IO pathway, we complemented the anterograde viral injections into the SC (Figure 1) with dual retrograde viral injections into the medial and lateral parts of the IO (Figure 2A) following our established protocol that allows accurate targeting with minimal spillage (Dorgans et al (2022)). Example 10x and 20x confocal microscopy scans of IO are shown for one mouse (out of 3) in Figure 2A1, A2. Here, tdTomato fluorescence was limited to the medial-most regions of the IO, while eGFP fluorescence was was seen in the more lateral parts, including the ventral fold of the PO. Figure 2 C1 is composed by summing the aligned image stacks (20x) acquired from 50 *µ*m sections, and shows that only a small fraction of the IO was double-labeled. Figure C1 also shows minimal labeling outside the IO, confirming that the labeled SC neurons specifically project to the IO. As shown in Figure 2C2, the relative extents of the GFP and tdTomato labeling were consistent among the 3 animals (tdTomato: 4.98 +- 0.60 %; eGFP: 6.95 +- 0.53 %) and the extent of double labeling was always minor (0.56 +- 0.16 %).

**Fig. 2.**
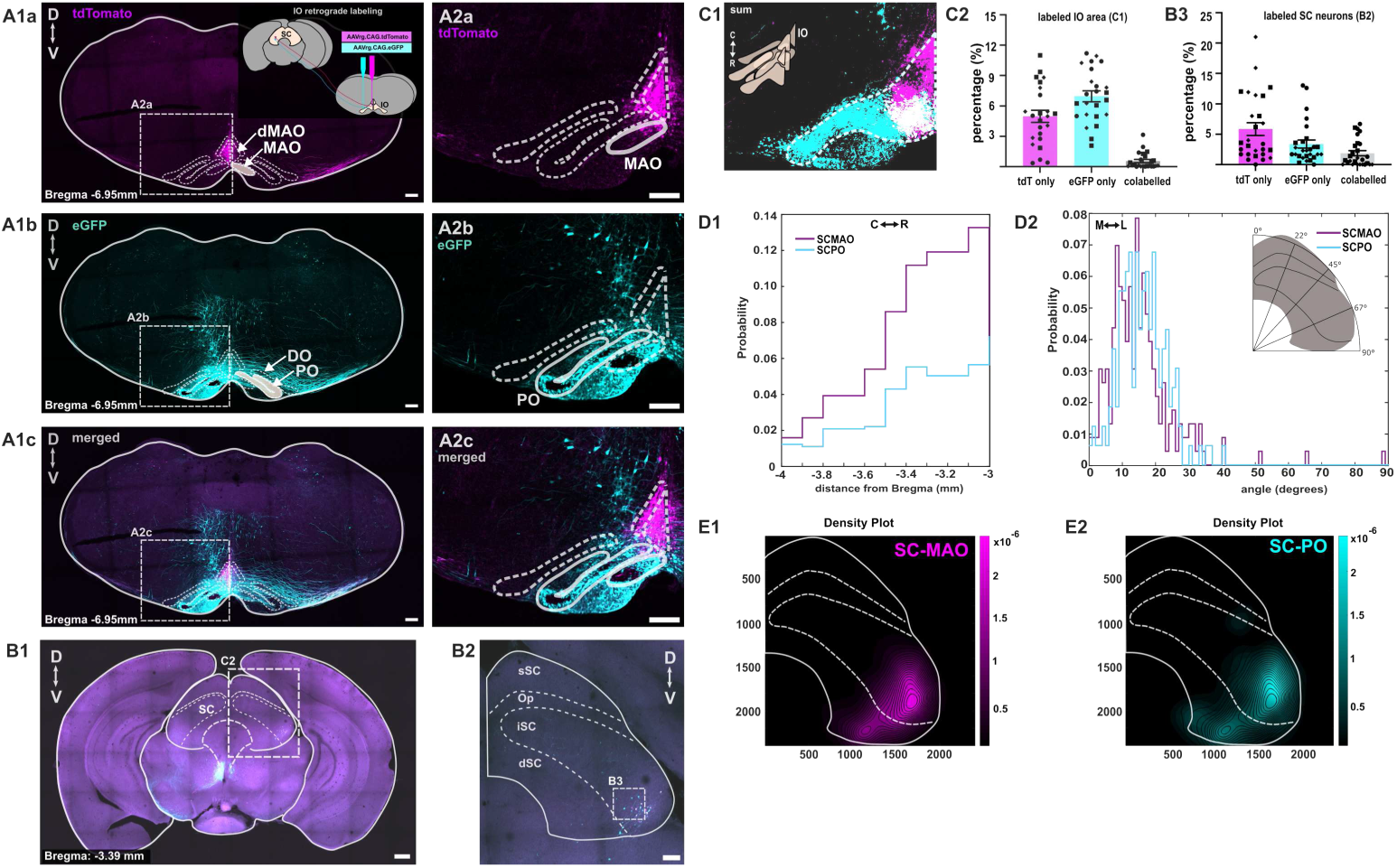
Identification of SC-MAO and SC-PO neurons through IO retrograde labeling. (**A1**) Schematic illustrating the retrograde labeling of different part of IO with AAVrg.CAG.eGFP labeling the lateral part of IO, including the PO and AAVrg.CAG.tdTomato labeling mainly restricted to MAO. (**A2**) Higher magnification of AAVrg injections into PO and MAO shown in A1. (**B1**) Coronal section of the midbrain with the superior colliculus and its retrogradely labeled neurons. (**B2**) Higher magnification of labeled tdTomato and eGFP neurons in the superior colliculus. (**C1**) Summation of labeled IO areas with tdTomato (SCMAO), eGFP (SCPO), and both, shown in panel A1-2 across the rostro-caudal axis (n=3). Percentage of this labeled IO area in C1 is shown in (**C2**). (**B3**) Percentage of SC neurons labeled with eGFP, tdTomato, and both across the rostro-caudal axis (n=3), indicated by the square in panel B2.(**D1**) Stairs histogram of labeled SCMAO and SCPO across the rostro-caudal axis, and (**D2**) based on the angular coordinates. (**E1-E2**) Density plot of SC neurons that exclusively project to MAO and PO. scalebar of 500 *µ*m for SC overview and scalebar of 200 *µ*m for the rest of the figures.

We then manually identified all neurons within the SC that were retrogradely labeled with tdTomato, eGFP, or both (Figure 2 B2, B3). The majority of the neurons expresed either tdTomato or GFP; overall, less than 5 % of them were double-labeled (from 27 SC slices from 3 animals). Notably, despite the fact that tdTomato-labeled IO regions covered less area than those labeled by GFP (Figure 2C2), majority of labeled somata within the SC expressed tdTomato, in line with the known strong projection from SC to MAO. However, a considerable number of SC neurons labeled with eGFP were always found intermingled with those labeled with tdTomato even when the retrograde virus expressing eGFP was not seen to extend near the MAO. As only a minority of neurons expressed both fluorophores in Figure 2B2-3 and 3B1), it seems that divergence of axons from SC neurons to both IO regions is not common.

**Fig. 3.**
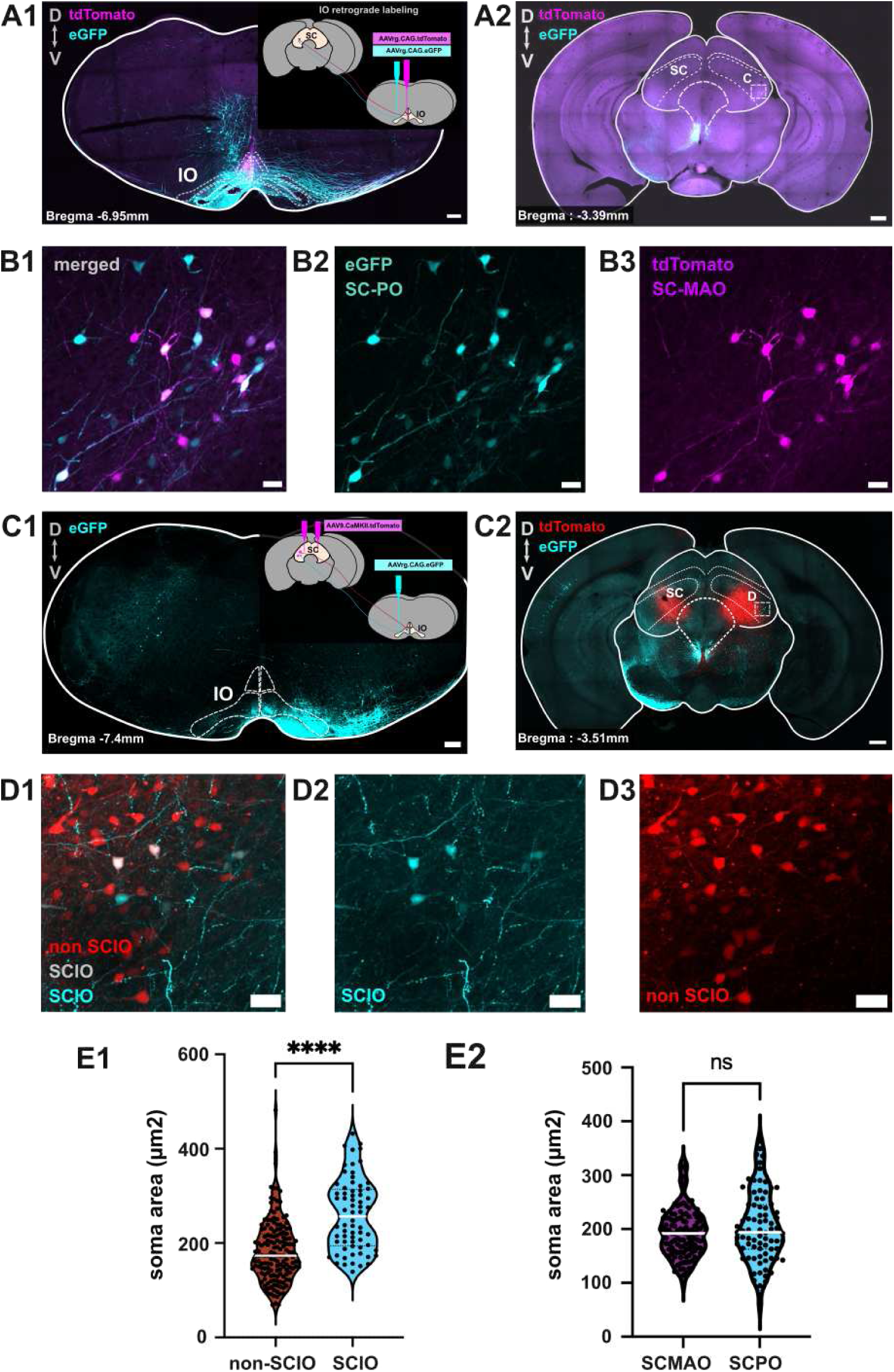
Comparison of SC-MAO, SC-PO and non IO projecting SC soma size. (**A1**) Schematic illustrating the retrograde labeling of different part of IO with AAVrg.CAG.eGFP labelling the lateral part of IO, including the PO and AAVrg.CAG.tdTomato labeling mainly restricted to MAO as shown in Figure02. Scalebar: 200 *µ*m (**A2**) Coronal section of the midbrain with the superior colliculus and its retrogradely labeled neurons, showing tdTomato, eGFP and colabeled neurons. Scalebar: 200 *µ*m (**B1-3**) a higher magnification of area indicated by square in the (**A2**), showing colabeled neurons, eGFP and tdTomto labeled neurons. Scalebar: 20 *µ*m (**C1**) Schematic illustrating the retrograde labeling of SC-IO neurons by injecting the AAVrg.CAG.eGFP into the IO and labeling of SC neurons with AAV9.CAMKII.tdTomato. Scalebar: 200 *µ*m (**C2**) Coronal section of the midbrain with the tdTomato labeled SC neurons. Scalebar: 200 *µ*m (**D1-3**) a higher magnification of area indicated by square in the (**C2**), showing SC-IO neurons and non-IO projecting SC neurons. SC-IO neurons are indicated by the eGFP and colabeled neurons while non SC-IO neurons indicated by tdTomato-only labeled neurons. Scalebar: 20 *µ*m (**E1**) Violin plot comparing the soma size between non SC-IO and SC-IO neurons. (**E2**) Violin plot comparing the soma size between non SC-MAO and SC-PO neurons. Mann-Whitney test was used to measure the statistical significance.

For a more quantitative examination of the localization of SC-IO neurons, we registered the labeled soma coordinates to a reference space (see Methods) to assess whether SC-MAO and SC-PO neurons occupied similar regions within the superior colliculus. Indeed, in all animals, the SC-MAO and SC-PO neurons were intermingled without evident differences for “angular” location (Figure 2 D2). Furthermore, no preferred rostro-caudal regions were identified for either SC-MAO and SC-PO neurons (Figure 2 D1) and density map found no preferred regions for either group (Figure 2 E1-2). However, we observed a higher density of SC-MAO and SC-PO neurons in the rostral SC compared to the caudal SC (Figure 2 D1). Thus, we conclude that the SC-MAO and SC-PO neurons occupy the same regions of the SC, despite their projections being exclusively targeted at different part of the IO.

### 2.3 Comparison of SC-MAO, SC-PO and non IO-projecting SC neurons

Since no differences were found in the soma locations of the SC-MAO and SC-PO neurons, we further investigated whether these neurons could be distinguished by their soma sizes. Using z-projection (standard deviation) images from 20x confocal stacks of labeled SC-MAO and SC-PO neurons (Figure 3A-B), we manually traced the labeled somata to measure their areas. The violin plot (Figure 3E2) did not reveal significant differences in the size of the soma (*P* = 0.45). However, SC-PO neurons exhibited greater variability in size (average size of SC-MAO: 193.7 ± 4.20 *µ* m-2;SC-PO: 203.4 ± 7.25 *µ*m-2).

We then compared the soma sizes between SC-IO and non-SC-IO neurons. To allow comparison between the IO-projecting and other SC neurons, we injected tdTomato-coupled AAV9 tracer under CaMKII promoter into the SC to reveal general neuronal shapes along with the eGFP-coupled retrograde viral tracer into the IO (Figure 3C). In these samples, we compared the sizes of eGFP expressing (IO-projecting) and non-expressing (non-projecting) neurons, and found that SC-IO neurons were larger than non-SC-IO neurons(*P* =*<* 0.0001, average SC-IO soma size 182.9 *µ*m2, average nonSC-IO soma size 259.8 *µ*m2) (Figure 3E1).

### 2.4 Immunohistochemistry of the SC-IO axons

Although it has long been presumed that most IO inputs that originate outside the cerebellar nuclei are primarily glutamatergic, recent findings have revealed additional complexities (Hoogstraten et al (2024)). We used immunohistochemistry to confirm the neurotransmitter identity of SC-MAO and SC-PO axons. SC neurons were anterogradely labeled (Figures 4A, 5A), followed by labeling of the vesicular glutamate transporter 2 (VGLUT2) and the vesicular *γ*-aminobutyric acid (GABA) transporter (VGAT) on SC-MAO (Figure 4) and SC-PO boutons (Figure 5). We then examined the colocalization of VGLUT2 and VGAT in labeled SC axonal boutons to identify the glutamatergic and GABAergic presynaptic boutons. As shown in Figures 4E and 5E, we found that less than 10% of SC-IO boutons colocalized VGLUT2 and VGAT, and the glutamatergic neurotransmitter type is indeed the major form. In MAO, most of the SC axons were VGLUT2 + (average: 92. 94%), followed by VGLUT2 + / VGAT + (average: 5. 48%) and VGAT + only (average: 1. 56%; Figure 5F). The percentage of VGLUT2 + axons in MAO was higher than in PO, although this difference was not statistically significant (*P* = 0.25). This variation may be due to the low percentage of VGLUT2+ boutons and a high percentage of VGAT+ boutons in one of the samples. In PO, most of the axons were VGLUT2 + (75. 62%), followed by VGAT + axons (21. 5%), which was higher than the percentage of VGAT + in SC-MAO, though not statistically significant (*P* = 0.1851). The percentage of VGLUT2+/VGAT+ axons in the PO was slightly lower than in the SC-MAO, at 2.9%.

**Fig. 4.**
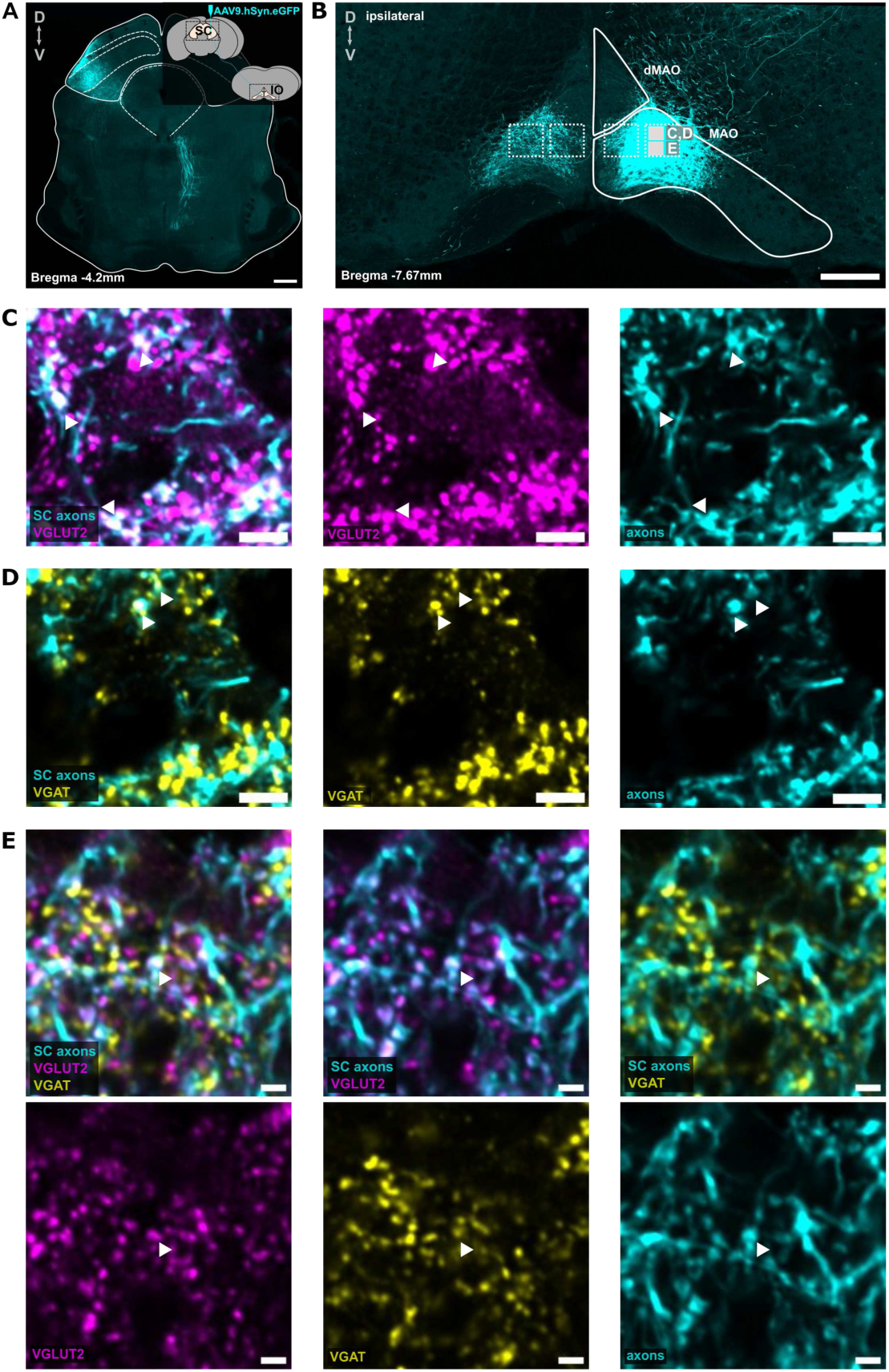
VGLUT2 and VGAT labeling in the medial accessory olive. (**A**) Anterograde labeling of SC neurons using AAV9.hSyn.eGFP, showing labeled SC axons in the medial accessory olive (MAO), as seen in (**B**). The 100×100 *µ*m squares indicate different regions of interest (ROIs) for 40x confocal acquisition of labeled VGLUT2, VGAT, and SC axons. (**C**) A 40X confocal acquisition of a section of the MAO area indicated by the square in panel B, displaying labeled SC axons and VGLUT2, SC axons and VGAT (**D**), and SC axons with both VGLUT2 and VGAT (**E**). White triangles in (**C-E**) indicate boutons colocalized with VGLUT2,VGAT, or both. The scalebar of 200 *µ*m for panel A,B and 5 *µ*m for panel C-E.

**Fig. 5.**
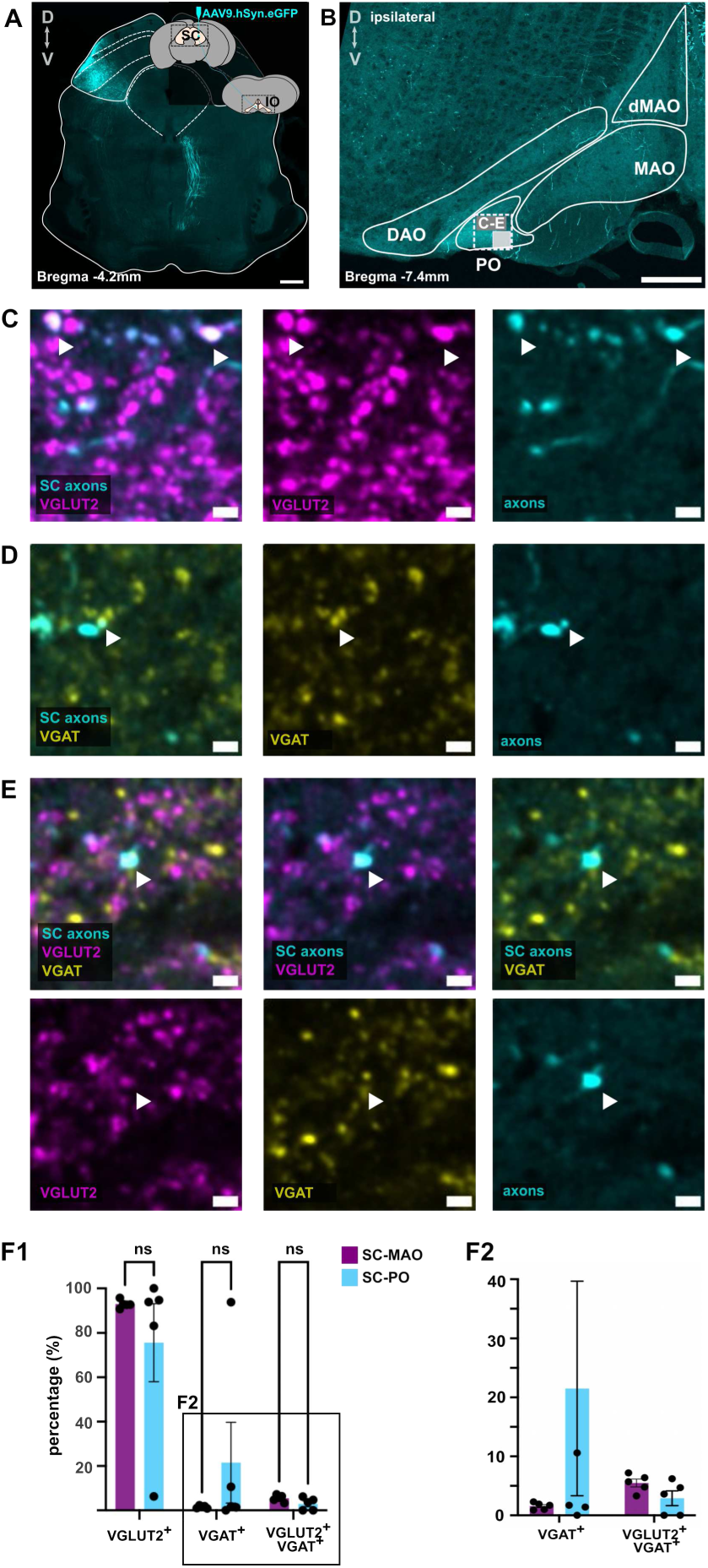
VGLUT2 and VGAT labeling in the principal olive. (**A**) Anterograde labeling of SC neurons using AAV9.hSyn.eGFP, showing labeled SC axons in the principal olive (PO), as depicted in (**B**). The 100×100 *µ*m squares highlight region of interest (ROI) for 40x confocal imaging of labeled VGLUT2, VGAT, and SC axons. (**C**) A 40x confocal acquisition of a section of the PO, indicated by the square in panel B, showing labeled SC axons and VGLUT2. (**D**) Confocal acquisition of the same section of PO showing labeled SC axons and VGAT, and (**E**) panels showing SC-PO axons colabeled with both VGLUT2 and VGAT (**E**) The white triangles indicate SC boutons that are collocalized with VGLUT2 and VGAT.(**F1**) Bar plot showing the percentage of SC axons that are VGLUT2+, VGAT+, VGLUT2+ and VGAT+ in MAO and PO. Dots represent data from five independent experiments. Multiple comparisons using two-way ANOVA is used to compare the percentage of VGLUT2+, VGAT+, and VGLUT2+/VGAT+ boutons in SCMAO and SCPO.(**F2**) The zoom in of VGAT+ and VGLUT2+/VGAT+ boutons.The scalebar of 200 *µ*m for panel A,B and 5 *µ*m for panel C-E.

### 2.5 Superior colliculus axons target the dendrites of the inferior olive neurons

Locating where SC ends within inferior olive neurons provides information on how SC input affects IO activity, given the dendritic gap junction coupling of IO neurons. Very few studies have examined subcellular axonal targeting patterns in IO (de Zeeuw et al (1989)), and the complex neuropil of IO, combined with the absence of clear layering in the IO, makes such investigations challenging.

We utilized our recently developed IO-specific virus that drives sparse labeling and anterogradely labeled the SC axons. 40X confocal images were acquired from the MAO (Figure 6C) and the PO (Figure 6D) regions to analyze putative synaptic contacts, where the labeled boutons and dendrites are in close proximity (*<* 0.1*µ*m). These putative synaptic contacts were observed in the dendritic shaft or in or near the dendritic spines.

**Fig. 6.**
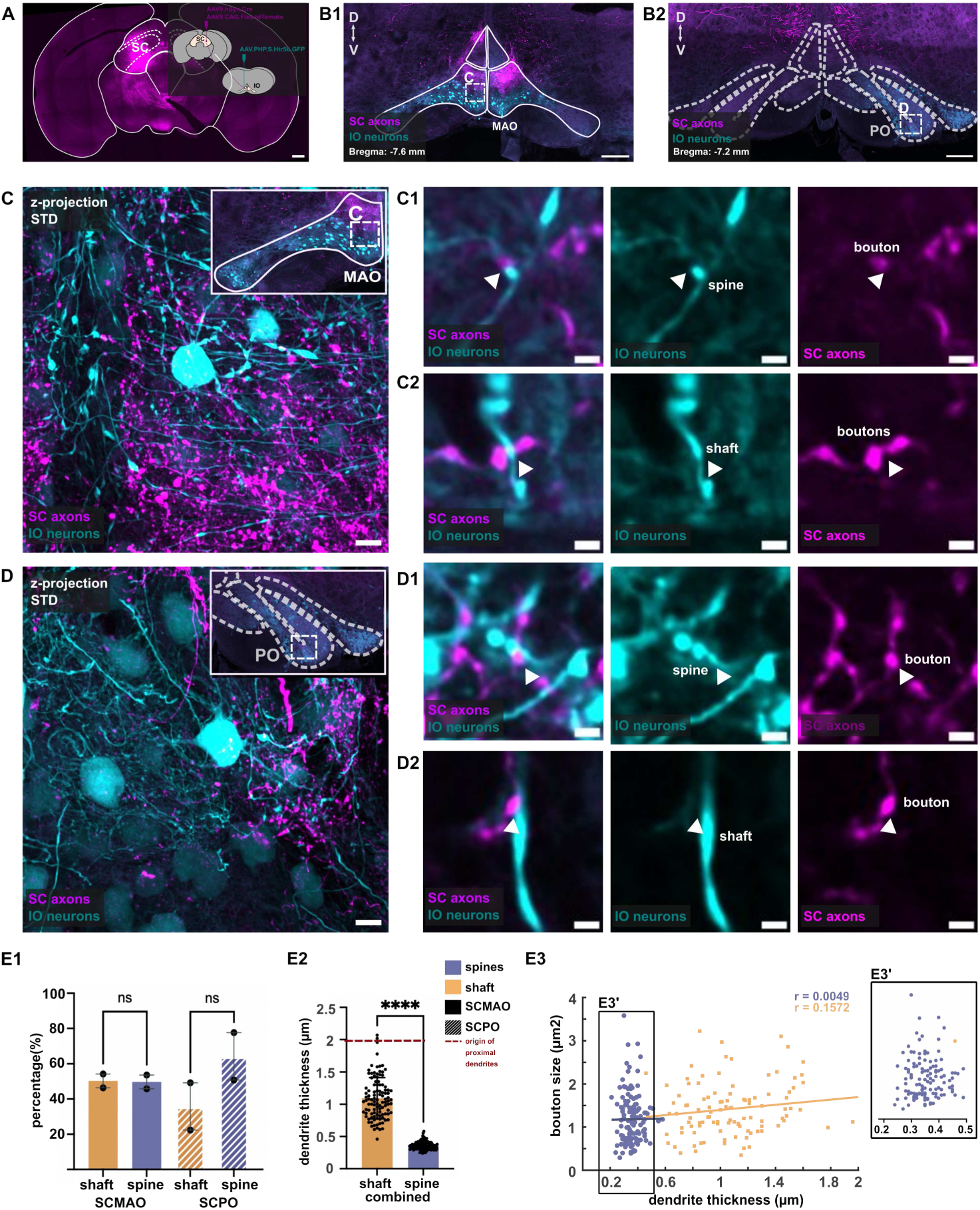
Identification of SC-IO synaptic contact along the SC dendrites. (**A**) viral tracer schematic to anterogradely label SC neurons using AAV9.hSyn.tdTomato and sparsely label the IO neurons using AAV.PHP.S.Htr5b.eGFP. (**B**) Inferior olive with tdTomato-labeled SC axons and eGFP-labeled IO neurons. Squares area in B indicates the ROI for 40X confocal imaging of the axons and IO neurons in MAO (**C**) and PO (**D**). (**C1, D1**) shows synaptic contact found around the spines in MAO and PO while (**C2, D2**) shows synaptic contact found around dendritic shaft in both regions. (**E1** Bar plot showing the number of synaptic contact on shaft and spines in SC-MAO and SC-PO. Mutiple comparison anova reveals that the higher number of spine contact in SCPO is not statistically significant.(**E2**) Bar plot showing the dendrite thickness of synaptic contacts found on the shaft and spines. A Mann-Whitney test revealed that the differences in dendrite thickness between shaft and spine synapses is statistically significant. (**E3**) Scatterplot showing the relationship between dendrite thickness and bouton size for shaft and spine contacts, with the Pearson correlation indicating a weak correlation between bouton size and dendrite thickness. (**E3’**) Zoomed-in view of the boxed area in (**E3**), highlighting the minimal overlap in dendrite thickness between shaft and spine synaptic

Of all boutons examined in SC-MAO (124 boutons, 2 animals), 50. 3% were located on dendritic shafts, while in SC-PO, 64. 2% of the boutons examined (123 boutons, 2 animals) were found in the dendritic spines (Figure 6E1). A two-way ANOVA did not reveal significant differences in the distribution of synaptic contact between the shaft and spine between SC-MAO and SC-PO (*P* = 0.32).

Since there were no differences in the distribution of the synaptic contacts of the spine and shaft between SC-MAO and SC-PO, we combined the data to examine the differences in dendritic thickness at the synaptic contacts of the shaft versus the spine (Figure 6E2). We observed significant differences in dendritic thickness, with spine contacts observed primarily in thinner dendrites (average: 0.36 ± 0.005 *µ*m) and shaft contacts occurring in thicker dendrites (average: 1.1 ± 0.03 *µ*m). This suggests that spine contacts are more likely to occur in thin and distal dendrites.

Furthermore, we analyzed the correlation between dendritic thickness and the size of the presynaptic boutons at the dendritic shaft and spine. Using Pearson’s correlation, we found a weak correlation between bouton size and dendritic thickness for both types of synaptic contacts (Figure 6E3). Figure 6 E3’ highlights the minimal overlap in dendritic thickness between the spine and the shaft synapses.

Lastly, our confocal data also showed that an axon can have two or more boutons on the same IO dendrite, which we observed in both MAO and PO. The 3D reconstruction of this observation is shown in Figure 7.

**Fig. 7.**
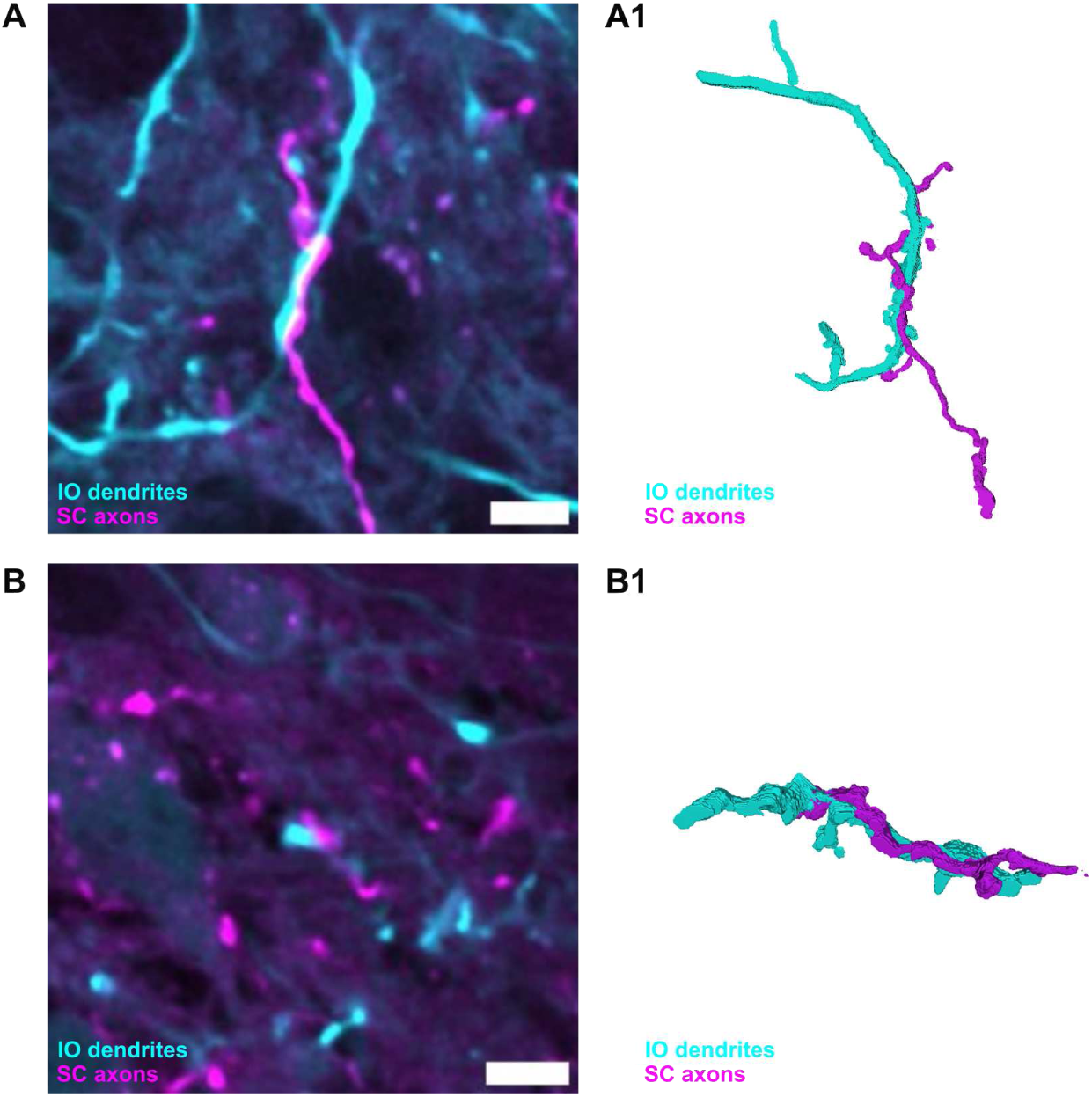
Reconstruction of SC-IO synapses. (**A**) Example confocal images of IO dendrites receiving two synaptic boutons from SC axon in caudal MAO and the PO dendrites. (**B**). (**A1, B1**) 2D visualization of 3D reconstruction of IO dendrites and SC axon from confocal image show in A, B.

### 2.6 In vivo effects of optogenetic activation of superior colliculus axons on the inferior olive activity

The presence of SC-IO synapses on dendritic spines raises the question of whether such synapses on distal dendrites are strong enough to evoke action potential in IO, or, would their function rather be related to gap junction coupling and network synchronization. To investigate these questions, we examined neuronal activity in the soma using a novel surgical methodology that allows calcium imaging and optogenetic activation of transfected axons in the ventral side of IO in living animals (Figure 8 A-D)Guo et al (2021)).

**Fig. 8.**
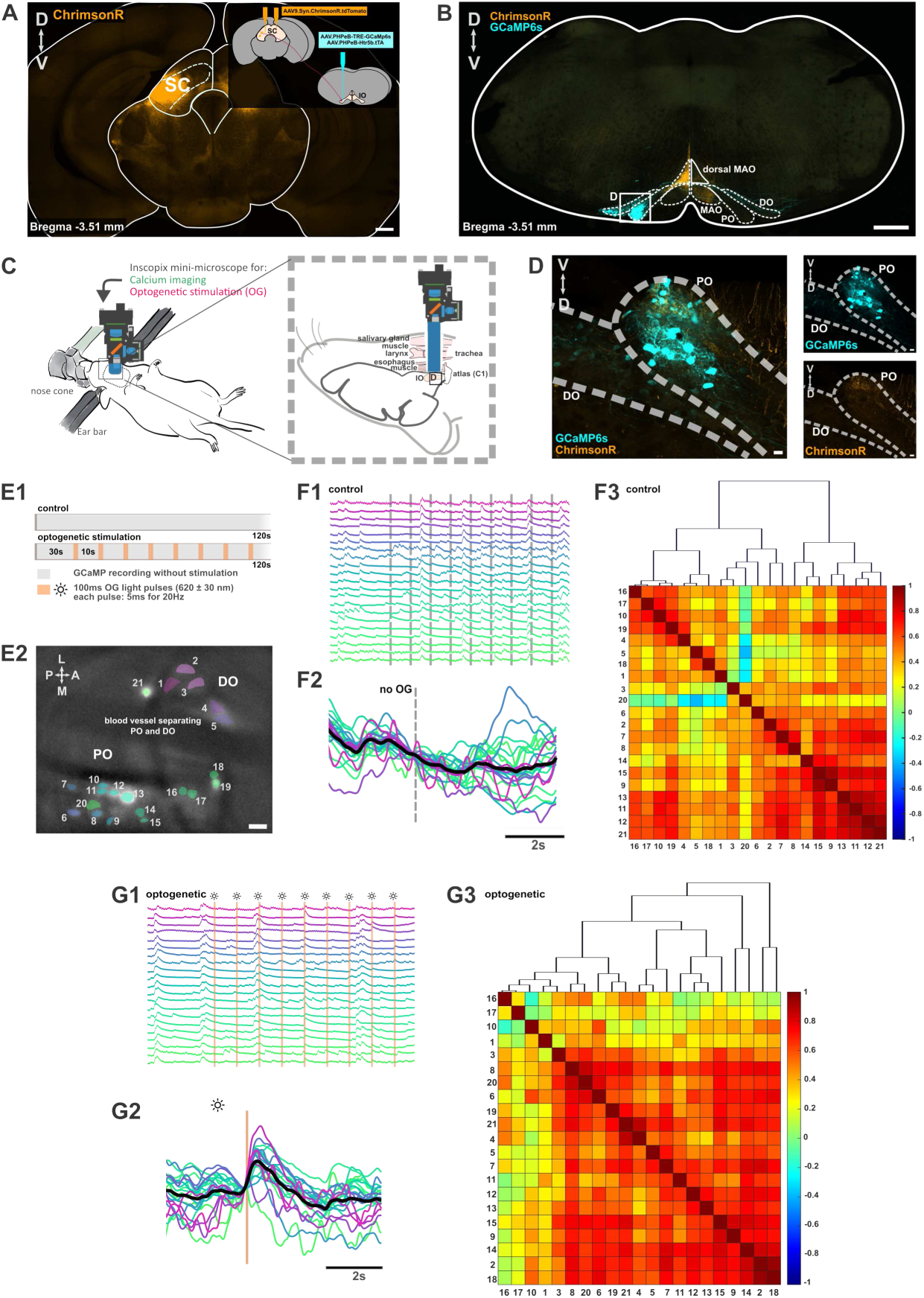
In-vivo calcium recording of IO neurons and the effect of optogenetic activation of SC axons on IO neurons activity. (**A**) Coronal section of midbrain containing SC neurons labeled with ChrimsonR coupled with tdTomato with a small schematic showing injection of AAV9.Syn.ChrimsonR.tdTomato into the SC and a mixture of AAV.PHPeB.TRE.GCaMP6s and AAV.PHPeB.Htr5b.tTA into the inferior olive. Scalebar 500 *µ*m (**B**) Coronal section of brainstem and the inferior olive with labeled SC axons and IO neurons. Scalebar 200 *µ*m (**C**) Schematic of ventral surgery to place the GRIN lens on the surface of ventral IO. (**D**) Enlarge view of IO indicated by the square box in B,C, showing labeled IO neurons and SC axons on the ventral leaf of the PO. Scalebar 20 *µ*m. (**E1**) Recording settings for control (GCaMP6s recording only) and optogenetic trials. (**E2**) Field of view of labeled and active IO neurons during recording with a scalebar of 20 *µ*m. The fluorescence traces of IO neurons on E2 during control (**F1**) and optogenetic trial (**G1**). The color of the traces corresponds the color of neurons in E2, F-G. (**F2, G2**) Summary of fluorescence traces across stimulations, plotted before, during and after stimulation. Each colors represents different neurons with average from all neurons is indicated with black trace. The dashed lines in F1 and F2 represent the timing of the stimulation events shown in G1 and G2, without the actual stimulation being applied. (**F3**) Cross correlation matrix and dendogram of recorded neurons during control and optogenetic trial (**G3**).

In these experiments, conducted in 2 animals, SC axons targeting the PO region were transfected with ChrimsonR and stimulated by brief pulses of light (10-second intervals; Figure 8 E). As shown in Figures 8 F1-2, the network shows a significant synchronization of activity even without stimulation (Figure 8 F3). In stimulation trials (Figure 8 G1-2), the light robustly evoked responses in majority of the cells.

We then examined whether light stimulation has any effect on network synchronization. To ensure that the syncrhonization effect extends beyond the direct stimulation effect, we exclude the traces during the stimulation-evoked events and examine the correlation matrix in control vs. optogenetic trial. The cross-correlation matrix shows an increase in synchronicity between IO neurons in the optogenetic trial (mean +- SEM: 0.5119 +- 0.0107, Figure 8 G3), compared to the control trial (mean +- SEM: 0.4646 +- 0.0109, Figure 8F3).

## 3 Discussion

In this study, we describe the projections between the mouse superior colliculus and the inferior olive in detail. In doing so, we provide evidence for a novel pathway linking the superior colliuclus to a region of the principal inferior olive. The SC axon terminals are largely glutamatergic and target distal dendritic spines and shafts. We also show that optogenetic activation of the SC terminals in the principal IO region that targets Crus 1 in the cortex can not only generate action potentials, but also lead to long-lasting synchronization of the olivary circuit. These observations suggest that the SC-IO projection is involved in motor coordination beyond posture and balancing. In the following, we will elaborate on the implications of findings and discuss future directions.

### Lateral part of the intermediate and deep layers of the superior colliculus sends mainly glutamatergic projections to the medial accessory olive and the principal olive

As the first step in examining subcellular targeting of SC axons, we anterogradely label SC neurons and confirm previous findings showing the projections from the medial and lateral part of the SC to distinct parts of the contralateral cdMAO, along with weaker projections to the ipsilateral side. To our surprise, we found additional SC projections in the ipsilateral ventral principal olive (PO) (Figure 1). This SC-PO projection was further validated through our retrograde labeling experiment (Figure 2). Examination of retrogradely labeled SCMAO and SCPO neurons revealed that they intermingle in the same region and layers of the lateral SC (Figure 2 B-E) and that they are very similar in size (SCMAO = 193.7±4.201*µ*^2^*, SCPO* = 203.4 ± 7.250*µ*m^2^)(Figure 3).

However, the distribution of SC-MAO and SC-PO neurons along the rostro-caudal axis of SC showed a higher number of SC-MAO neurons compared to SC-PO neurons (Figure 2 D1), which may explain the lower probability (30-35%) of detecting SC-PO axonal projections (Figure 1H2). Examination of the distribution of the presynaptic VGLUT2 + and VGAT + boutons revealed no significant differences, and the majority of boutons were VGLUT2 + (Figure 4,5).

The superior colliculus is well known for its role in mediating orienting-related movements (Masullo et al (2019); Zhou et al (2023); Solié et al (2022), and the SC-MAO pathway supports this function. MAO targets Purkinje neurons in the cerebellar vermis, which is essential for postural control and balance essential for orienting movements. Although the SC-MAO pathway aligns with the established role of SC in orienting movement, the projection of SC-PO to Purkinje neurons in CrusI areas suggests that this pathway might carry distinct information. As Crus I is related to orofacial movements and SC-IO neurons reside in the lateral SC, which processes signals from the lower visual field (Cang et al (2018); Ito and Feldheim (2018)), the SC-PO pathway could be involved in hunting-related behaviors.

### Superior colliculus axons target the dendrites of IO neurons

IO neurons are electrically coupled via gap junctions that are located mostly in dendritic spines (De Zeeuw et al (1998); Devor and Yarom (2002); De Gruijl et al (2014)). This gap junction coupling enables neurons to share electrical activity and is proposed to mediate synchronization within the IO population (Leznik and Llinás (2005)). GABAergic input near the gap junctions is known to disrupt this synchronization by acting as an electrical shunt (Lefler et al (2014). Since SC-IO boutons are predominantly glutamatergic, as indicated by the large proportion of VGLUT2 + SC boutons, input from the SC may evoke IO spikes. However, in IO, the effect of presynaptic boutons could also depend on their location along the IO neurons. If the presynaptic boutons are located primarily on the distal dendrite, there is the possibility that the SC input influences the synchronization of the neuronal population. Therefore, we examined the subcellular localization of SC-IO synapses and investigate if there are differences between SC-MAO and SC-PO.

We identified SC boutons and IO neurons that are extremely close to each other (*<* 0.1 *µ*) or slightly overlapped at their edges. We found boutons mainly on dendrites, as we were unable to confirm the presence of somatic contact. Along the dendrites, we found putative contact on the shaft or dendritic spines, where most of the gap junctions are located. We found no preferences for shaft or spine contact, but dendritic thickness analysis shows that shaft and spine contact is found in different parts of the dendrites. Synapses on the shaft are found on thick dendrites closer to the soma, known as proximal dendrites, whereas synapses on the spines are mostly found in thin dendrites further away from the soma, known as distal dendrites. This finding confirms the hypothesis that most of the dendritic spines of the IO are located in the distal dendrites. We also found that an axon can have multiple synaptic boutons contacting the same location on a dendrite, which can amplify the strength of the synaptic input. Furthermore, the presence of glutamatergic synaptic contact in the dendritic spines suggests the two possible effects of the SC-IO pathway: (i) SC activation increases the probability of spiking in IO neurons and/or (ii) SC activation influences synchronization among the IO population. To test these possibilities, we optogenetically activated SC boutons and examined the effect on the calcium activity of the IO in an anesthetized mouse.

### Optogenetic activation of superior colliculus axons drives calcium events in inferior olive neuronal population in-vivo

Our findings demonstrate that SC-PO synapses can increase the probability of spikes and synchrony within the IO population. However, the SC input does not reliably evoke a calcium event (spikes) in IO. Full trial traces in figure 8G1 indicate that optogenetic activation of the SC boutons evokes spikes in approximately 1 out of every 3 stimulation events, on average. Despite the synapses being distal to the soma, the SC input is strong enough to evoke spikes in the IO. This strength may be attributed to the presence of multiple synapses from a single axon, which could amplify synaptic input and facilitate spike generation in IO and complex spikes in the Crus I Purkinje neurons. This finding aligns with previous studies showing that SC activation induces complex spikes in the vermis (Akaike (1985), which receives climbing fiber projections from the medial accessory olive. There is also the possibility of somatic synapses being present, although this cannot be confirmed with the current data.

Previous study showed that GABAergic input near the gap junctions can decrease electrical coupling and synchronicity between neurons (Lefler et al (2014)). Here, we show that optogenetic activation of glutamatergic presynaptic SC boutons increases synchrony among the neuronal population of IO (Figure 8 G3). This finding aligns with our anatomical observation of SC-IO synapses on dendritic spines, where the gap junctions are located. IO sends climbing fibers that innervate several (± 6) Purkinje neurons in the cerebellum (Sugihara et al (2001)). Increased neuronal synchrony suggests that more Purkinje neurons receive information simultaneously, potentially enhancing coordinated cerebellar output.

We noted the presence of labeled SC axons in the dorsal region of the medial accessory olive (encompassing the beta subnucleus, the cap of Kooy, and the ventro-lateral protrusion), which were absent from our initial labeling of the SC-PO pathway. We suspect that this labeling may result from the diffusion of anterograde tracers into adjacent regions, such as the dorsal periaqueductal gray (PAG) or the inferior colliculus (IC).

## 4 Conclusion and Future Directions

Our study provides a mesoscale and microscale description of the SC-IO pathway, including the novel SC projection to the PO. We showed that the pathway is mainly glutamatergic with putative synapses found on the shaft of the proximal dendrite and on the spines of the distal dendrites. We further showed that optogenetic activation of the SC boutons can evoke spikes and increase the synchrony among the IO neuronal population. While the SC-MAO pathway is known to play a role in postural control and balance, the function of the SC-PO pathway is less understood. Its projection to Crus I suggests potential involvement in cognition or orofacial movement. Given that most SC neurons involved in this pathway are located in the lateral SC, representing the lower visual field, we hypothesize that the SC-PO pathway may play a role in hunting-related behaviors. Future experiments comparing the effects of SC-PO and SC-MAO activation or inhibition will be important to understand the role of the olivocerebellar system in orienting related movement.

## 5 Methods

### Animals

In the current study, wild-type C57BL / 6J male mice (CLEA Japan, Shizuoka, Japan) were used with an age of approximately 12-16 weeks. The animal experiment was conducted in strict adherence to the approved guidelines by the Okinawa Institute of Science and Technology (OIST) and the Institutional Animal Care and Use Committee (IACUC), within a facility accredited by the Association for Assessment and Accreditation of Laboratory Animal Care (AAALAC International). Of these animals, 4 mice were used for anterograde labeling of SC neurons, 3 mice were used for retrograde labeling of IO afferents, 3 mice were used to label SC axons and sparsely label the IO neurons, and another 3 mice of mice were used for anterograde labeling of SC neurons combined with immunofluorescece. 2 mice were used for in-vivo calcium imaging experiment.

### Stereotaxic surgery

Prior to the surgery, the mice were anesthetized with 5pct of isofluorane (SomnoSuite, Kent Scientific, CT, USA) and placed on the stereotaxic frame equipped with a heat pad (38°C; TMP-5b, Supertech Instruments). During surgery, the animal was kept under anesthesia with 1.5 - 2. 5% isofluorane through the nose cone. Precise alignment of the head and body is crucial for accurate targeting of deep structures such as the IO (Dorgans et al., 2020; Guo et al., 2021). After scalp shaving, local anesthesia was administered using Xylocaine gel (Xylogel, gel 2pct, Aspen, Japan), and the skin was disinfected and incised with a scissor to expose the skull. Subsequently, small craniotomies (approximately 1 mm in diameter) were created using a handheld drill (Surgic XT Plus drill, NSK Dental, Japan).

Following the placement of viral mixtures into the quartz capillary pipette, the pipette was gently maneuvered into the desired location within the brain tissue at a slow speed of approximately 0.2 mm/s. Using bregma as a reference point, the coordinates of AP: 3.3mm, ML: 1.2mm, DV: 2.2mm and AP: 3.5mm, ML: 1.4mm, DV: 2.4mm were used to target the superior colliculus neurons. The coordinates of AP: 6.0mm, ML: 0.4mm, DV: 6.75mm and AP: 6.4mm, ML: 0.2mm, DV: 6.75mm were used to target the PO and MAO consequently. At these locations, 20-40 nl of viral mixtures were injected with a speed of 20 nl / min. Approximately 5-10 minutes after the completion of the viral delivery, the pipette was retrieved from the tissue with slow speed. The skin was clean and the incision was sutured, followed by subcutaneous administration of 5mg/kg of Rymadil (Zoetis, New Jersey, US).

### Tracers

For anterograde labeling of superior colliculus neurons (SC), we use AAV9.hSyn.eGFP (final titer: 7 × 10^11^ vg / ml, 50465, Addgene) and a mixture of AAV9.hSyn.Cre (105553, Addgene) or AAV9.CamKII.Cre (105558, Addgene) with AAV9.CAG.Flex.tdTomato (28306, Addgene). These viral vectors were carefully mixed and subsequently diluted in phosphate buffered saline (PBS) to achieve a final titer of 1.7 × 10^12^ vg / ml.

To retrogradely label the afferents in different inferior olive subnuclei (IO), we used AAVrg.CAG.tdTomato (final titer: 3.5 × 10^12^ vg/mL, 59462, Addgene) and AAVrg.CAG.eGFP (final titer: 3.5 × 10^12^ vg / mL, 37825, Addgene) in the medial accessory olive (MAO) and principal olive (PO).

The AAV9.hSyn.Cre and AAV9.CAG.Flex.tdTomato mixture, along with AAV.PHP.S.Htr5b.eGFP (final titer: 5 × 10^12^ vg/mL, OIST), was used to label SC axons and sparsely label IO neurons. For *in vivo* calcium imaging of IO neurons, we used a mixture of IO-specific viruses: AAV.PHP.eB.TRE.GCaMP6s (final titer: 2 × 10^11^ vg/mL, OIST) and AAV.PHP.eB.Htr5b.tTA (final titer: 2 × 10^10^ vg/mL, OIST) Dorgans et al (2022). Furthermore, AAV9.Syn.ChrimsonR.tdTomato was used for optogenetic activation of SC axons.

### Tissue Processing and Immunohistochemistry

Two to three weeks after injection of viral tracers into the superior colliculus (SC), the animals were transcardically perfused with PBS followed by 4% PFA (w/v, in PBS). The brains were extracted and postfixed in 4% PFA solution for 2-3 hours at room temperature, after which they were stored in PBS at 4 ° C until sectioning. The brains were sectioned using a vibratome (Model 5100MZ-plus, Campden Instruments) equipped with ceramic blades (Model 7550-1-C, 38 x 7 x 0.5 mm, Campden Instruments). The slices were mounted with vectashild (vector laboratories) or prolong glass (invitrogen) and #1.5 coverslip glass (Harvard Apparatus, MA). For immunohistochemistry, the brains were sectioned with vibratome or using a cryostat (CM1950, Leica) equipped with blades (Disposable Blades DB80LX, Leica). The brains that were sectioned with the cryostat were cryoprotected by submerging them in a 10% sucrose solution (w/v, in PBS) for several hours at room temperature, followed by being stored overnight in a 30% sucrose solution (w / v, in PBS) before sectioning. To verify the injection sites in the SC, the rostral parts of the brains were sectioned coronally at 50 *µ*m or 100 *µ*m. These sections were washed 2x in PBS for 10 minutes at room temperature, followed by 3 × 5 minute washes in 0.1M phosphate buffer (PB) at room temperature, after which they were mounted on an objective glass with Prolong glass mounting medium (invitrogen) and #1.5 coverslip glass (Harvard Apparatus, MA).

The brainstem were sectioned separately at 50 *µ*m in coronal orientation, also using the vibratome with ceramic blades or cryostat. For immunostaining, one out of two brainstem sections was stained freely floating using anti-VGluT2 (Polyclonal, Thermofisher) and anti-VGAT (Polyclonal, Alomone labs). The sections were first washed 4 times in PBS at room temperature for at least 10 minutes. Subsequently, they were blocked and permeabilized in PBS containing 10% Normal Goat Serum (NGS, Abcam) and 0.5% Triton X (Thermofisher.) for 2 hours on a shaker (100 rpm), followed by primary antibody incubation. For this step, the slices were incubated for 48 to 72 hours at 4°C in a solution of PBS with 2% NGS and 0.1% Triton X and anti-VGluT2 (1:500) and anti-VGAT (1:2000) primary antibodies. The sections were subsequently washed 5 times for 10 minutes in PBS at room temperature, before secondary antibody incubation for 2 hours at room temperature, in a PBS solution with 2% NGS, 0.1% Triton X and secondary antibodies. The secondary antibodies used were goat-anti rabbit-A647 (Invitrogen) and goat-anti guinea pig-A555 (Invitrogen), both at a concentration of 1:2000. Finally, the sections were washed 2 times for 10 minutes in PBS, followed by 3 washes in 0.1M phosphate buffer (PB), before being mounted using the same procedure as the SC sections.

### Confocal acquisition and Image processing

The z-stacks images of the sections labeled with viral and immunohistological methods were acquired using a Zeiss LSM 880 confocal system (Zeiss, Germany). For the overview scanning of entire cortex and brainstem, 10X objective (Plan-Apochromat 10x, NA, Zeiss, Germany) with 6*µ*m z-step were used. To scan the IO region, 20x objective Plan-Apochromat 20x, NA, Zeiss, Germany) with 3-6*µ*m z-step were used. To scan the entire SC region, 10x objective Plan-Apochromat 20x, NA, Zeiss, Germany) with 6*µ*m z-step were used. For higher magnification images, 40x objective (Objective “Plan-Apochromat” 40x Oil DIC M27; NA 1.4; using Zeiss Immersoil oil; Zeiss, Germany) with around 0.07-0.1*µ*m zsteps was used, the value was adjusted to match the x and y of the scanned image. All images were acquired with a total image size of 1024 x 1024 pixels. For multichannel images, different excitation/emission wavelengths were used. Argon 488nm / 490–535 nm were used for eGFP, 561 nm / 470–655 nm for tdTomato and 633 nm / 638-758 nm for infrared. Different acquisition parameters were used for different experiments, but were kept constant in different samples within the experiments. After acquisition, the stacks images were processed by applying a Gaussian filter 3D with value of 1 for the x, y and z dimension and z projection was applied to acquired 2D image. For vizualisation and analysis, we used the standard deviation for the z projection and adjusted the histogram accordingly.

### Percentage of SC-IO axons labeling across IO subnuclei

After confocal acquisition and image processing, We categorized IO into three regions: caudal IO, middle IO, and rostral IO. The subnuclei within each IO region were manually delineated using Fiji software, and the intensity values of labeled SC axons in each subnucleus were measured across slices. In our analysis, we classified IOA, IOB, and IOC into ventral MAO and classified IOVL, IOBe, IOK, and IODM into dorsal MAO.

For each IO region (caudal, middle and rostral), we quantified the intensity of labeled SC axons in different IO subnuclei using multiple slices from five animals. Intensity values were normalized to the maximum intensity value for each animal (see supplementary Table 1). The average normalized intensity value in each subnucleus was presented in Figure 1H1. Furthermore, Normalized values below 0.2 were considered indicative of passing axons or background noise from image acquisition. Values above 0.2 were classified as 1 (presence of labeled axons), while values below 0.2 were classified as 0 (absence of labeled axons).

The percentage of subnuclei with labeled axons in the caudal, middle, and rostral IO regions is shown in Figure 1H2.

### Morphological measurement of SC-IO neurons

After image processing, all detectable SC-PO and SC-MAO neurons were delineated using Fiji software, subsequently saving them as Regions of Interest (ROIs). The soma with an unidentifiable shape or very faint fluorescent expressions is not counted as ROI. These ROIs were classified as SCMAO or SCPO neurons or both for the colabeled neurons and then measured for their area. The results were inputted into Prism for visualization and statistical analysis.

### Coordinate profile of SC-IO neurons

To locate the labeled SC-IO neurons along the rostro-caudal axis, the midline position of the acquired confocal images of the midbrain was manually aligned and the coordinates of each slice were matched to the bregma using the Mouse Brain Atlas (Paxinos & Franklin). Using Fiji software, all detectable SC-PO and SC-MAO neurons were manually traced, saved as ROIs, and the center of mass, X-coordinate of the midline, and Y-coordinate of the SC surface were measured. The x-y positions of the labeled SC-PO and SC-MAO neurons were normalized to the midline and SC surface. This x-y coordinate analysis was performed across slices and between animals. To further map the labeled neurons along the medio-lateral axis (M-L), the soma locations were recroded in “angular space”, following the conventions described in Benavidez et al (2021). The coordinate is shown in Figure 2D1-D2.

### Neuronal tracing and identification of the SC-IO putative synaptic contact

In the sections containing GFP+ SC axons in the inferior olive (IO), we identified several 100 x 100 m regions of interest within the caudal medial accessory olive (IOC) and principal olive (PO). In these ROIs, we quantified the ratio of VGluT2 + to VGAT + boutons in SC fibers. We purposely selected two ROIs within the IOC and one ROI in the ventral PO. These regions were scanned ipsilaterally and contralaterally to the injection site, totaling six ROIs per animal.

Z-stacks of these ROIs were acquired using a confocal microscope with a 40x objective. The images are further divided into four subregions to facilitate analysis. In each of the subregions, we applied noise reduction using the 3D Gaussian filter and manually quantified the percentage of GFP+ boutons labeled with VGluT2, VGAT, both and none, using the following procedures in the FIJI software. Afterwards, the analysis was performed in two steps using custom FIJI macros.

The first step aimed to identify potential GFP+ boutons in the image. For the subregions of the PO scans, we visually determined that the number of boutons was rather low, allowing us to identify and label all boutons in the image. This was done by manually drawing ROIs in FIJI using the free-hand or polygon selection tool. For the boutons in IOC,due to the high density of boutons, it was not feasible to label all boutons individually. Instead, a random subset of boutons was selected. This selection process was performed manually with the assistance of a macro. The macro identified local maxima that exceeded an intensity threshold of 2500 using a local maximum filter from the ImageJ 3DSuite package (Ollion et al (2013)). The macro then randomly presented to the person analyzing (observer) with one of the identified local maxima, allowing them to draw an ROI if the local maximum was deemed to correspond to a bouton. This process continued until all local maxima in the image were evaluated or a predetermined threshold (set at 50 boutons) was reached for the subregion.

The second step aimed to determine whether the boutons identified in step one were labeled with VGluT2 or VGAT. A macro presented each ROI to the user for manual categorization as labeled, unlabeled, or doubtful for both the VGluT2 and VGAT channels. For analysis purposes, boutons marked as “doubtful” were treated as unlabeled. Additionally, users could exclude any bouton from the analysis if they were not fully confident that the ROI represented a synaptic bouton.

The results were recorded and summarized in a Microsoft Excel file. To ensure reliability, each image was analyzed by two independent observers. For example, if step one was conducted by observer A, step two was performed by observer B.

### In-vivo calcium recording of IO neuronal population and Optogenetic activation of SC axons

Two to three weeks after injecting AAV9.Syn.ChrimsonR.tdTomato into the superior colliculus (SC) and AAV.PHPeB.TRE.GCaMP6s along with AAV.PHPeB.Htr5b.tTA into the inferior olive (IO), mice were anesthetized with 5% isoflurane for ventral surgery to allow placement of the GRIN lens (Guo et al (2021)). Prior to surgery, the thigh and throat were shaved, with thigh sensors used to monitor oxygen and breathing rate, and a rectal probe used to monitor body temperature.

An incision was made on the throat after the application of lidocaine. The salivary glands were displaced and the sternothyroid muscles overlying the trachea were removed for tracheotomy. The thyroid gland was removed and a thread was secured around the tracheal ring to stabilize it before connecting it to a pre-made intubation tube, ensuring the maintenance of the anesthesia. The esophagus, larynx, and surrounding tissues were removed, exposing the occipital bones, ventral arch, and anterior tubercle of the atlas. The ventral arches of the atlas and anterior tubercle were removed using a rongeur, and the surrounding occipital bone was cut to enlarge the field of view. The cartilage overlying the dura was removed to expose the ventral IO to have a clear view during calcium recording.

A GRIN lens (9 mm length, 1 mm diameter) connected to an Inscopix mini-microscope was placed on the ventral IO surface. For the GCaMP6s recording, blue LED light (455 ± 8 nm) was used, while red light (620 ± 30 nm) was used for optogenetics. The recordings were taken at 40 Hz for 120 seconds. During optogenetic stimulation, a 100-ms pulse train (20 Hz, 5-ms pulses) was delivered 30 seconds after the recording began, with nine repetitions with a 10-second interval between the stimulations. During recording, we reduced isofluorane and monitored the breathing rate to ensure that the animal was in the anesthesia state.

The recordings were spatially filtered and motion corrected using Inscopix Data Processing (IDP) software. The regions of interest (ROIs) were manually traced, and the fluorescence traces were analyzed and visualized in MATLAB.

## Supplementary information

### Supplementary Table

Table 1: The fluoresecence intensity of SC-IO axons in different IO subnuclei across rostro-caudal axis in 5 animals. The values were normalized to the highest intensity in each animals.

## Acknowledgements.

The authors would like to thank Hugo Hoedemaker and Dr. Guo Da for their assistance with the calcium recording experiment and Lakshmipriya Swaminathan for assisting with the analysis coding. The authors are grateful for the help and support provided by the Animal Resources Section (ARS) of Core Facilities at Okinawa Institute of Science and Technology Graduate University.

## Declarations

### Funding

This work was supported by Okinawa Institute of Science and Technology Intramural funding.

### Conflict of interest/Competing interests

The authors declare no conflict of interest.

### Author Contributions

DD: Contributing to experimental design, conducting experiments, data analysis, visualization, and manuscript preparation.

HN: Contributing to immunohistochemistry-related work such as experimental design, conducting experiments, data analysis, and manuscript writing.

MYU: Providing supervision and guidance in experimental design, data analysis, and manuscript preparation.

## Notes

### Competing Interest Statement

The authors have declared no competing interest.

