## Supplemental Table 1 for "Anatomical and functional examination of superior colliculus projections to the inferior olivary neurons in mice"

| | | MAO contra | MAO ipsi | dorsalMAO contra | dorsalMAO ipsi | IOPr contra | IOPr ipsi | IOD contra | IOD ipsi |
|---|---|---|---|---|---|---|---|---|---|
| caudal IO | SC080 | 1 | 0.1 | 0.7 | 0.1 | | | | |
| | | 1 | 0 | 0.3 | 0 | | | | |
| | | 1 | 0.1 | 0.7 | 0.1 | | | | |
| | | 1 | 0 | 0.3 | 0 | | | | |
| | | 1 | 0.1 | 0.7 | 0.1 | | | | |
| | | 0.8 | 0 | 0.2 | 0 | | | | |
| | SC081 | 0.5 | 0 | 0.2 | 0 | | | | |
| | | 0.5 | 0.1 | 0.3 | 0.1 | | | | |
| | | 1 | 0.1 | 0.4 | 0.1 | | | | |
| | | 1 | 0.1 | 0.6 | 0.1 | | | | |
| | | 0.9 | 0.1 | 0.3 | 0.1 | | | | |
| | | 0.9 | 0.1 | 0.5 | 0.1 | | | | |
| | SC010 | 0.3 | 0 | 0.2 | 0 | | | | |
| | | 1 | 0.1 | 0.3 | 0 | | | | |
| | SC011 | 1 | 0 | 0.3 | 0 | | | | |
| | | 0.9 | 0.1 | 0.2 | 0 | | | | |
| | | 1 | 0 | 0.3 | 0 | | | | |
| | SC014 | 0.9 | 0 | 0.3 | 0 | | | | |
| | | 0.9 | 0.1 | 0.4 | 0.1 | | | | |
| middle IO | SC080 | 0.9 | 0.8 | 0.19 | 0.18 | 0.23 | 0.12 | 0.51 | 0.51 |
| | | 0.42 | 0.2 | 0.05 | 0.05 | 0.05 | 0.03 | 0.14 | 0.14 |
| | | 0.97 | 0.98 | 0.33 | 0.26 | 0.12 | 0.11 | 0.48 | 0.48 |
| | | 0.27 | 0.23 | 0.1 | 0.07 | 0.02 | 0.02 | 0.11 | 0.11 |
| | SC081 | 0.25 | 0.19 | 0.05 | 0.04 | 0.02 | 0.02 | 0.12 | 0.12 |
| | | 0.34 | 0.31 | 0.07 | 0.05 | 0.04 | 0.04 | 0.19 | 0.19 |
| | SCIHC010 | 0.76 | 0.3 | 0.07 | 0.06 | 0.03 | 0.02 | 0.13 | 0.13 |
| | | 0.66 | 0.42 | 0.12 | 0.08 | 0.06 | 0.06 | 0.27 | 0.27 |
| | SCIHC011 | 0.35 | 0.17 | 0.03 | 0.03 | 0.03 | 0.02 | 0.13 | 0.13 |
| | | 0.17 | 0.17 | 0.06 | 0.04 | 0.04 | 0.04 | 0.1 | 0.1 |
| | SCIHC014 | 0.31 | 0.23 | 0.03 | 0.03 | 0.02 | 0.03 | 0.1 | 0.1 |
| | | 0.15 | 0.18 | 0.07 | 0.07 | 0.05 | 0.05 | 0.12 | 0.12 |
| rostral IO | SC080 | 0.18 | 0.19 | 0.23 | 0.17 | 0.52 | 0.61 | 0.35 | 0.35 |
| | | 0.04 | 0.04 | 0.05 | 0.04 | 0.11 | 0.14 | 0.08 | 0.08 |
| | | 0.27 | 0.26 | 0.22 | 0.2 | 0.71 | 0.78 | 0.49 | 0.49 |
| | | 0.06 | 0.06 | 0.05 | 0.05 | 0.15 | 0.18 | 0.1 | 0.1 |
| | SC081 | 0.17 | 0.19 | 0.13 | 0.1 | 0.3 | 0.45 | 0.25 | 0.25 |
| | | 0.3 | 0.34 | 0.22 | 0.16 | 0.55 | 0.52 | 0.45 | 0.45 |
| | | 0.1 | 0.09 | 0.08 | 0.07 | 0.19 | 0.31 | 0.18 | 0.18 |
| | | 0.18 | 0.15 | 0.13 | 0.11 | 0.33 | 0.37 | 0.32 | 0.32 |
| | | 0.05 | 0.05 | 0.05 | 0.04 | 0.15 | 0.33 | 0.1 | 0.1 |
| | | 0.09 | 0.09 | 0.09 | 0.07 | 0.26 | 0.29 | 0.19 | 0.19 |
| | SCIHC010 | 0.11 | 0.13 | 0.09 | 0.09 | 0.28 | 0.33 | 0.18 | 0.18 |
| | | 0.06 | 0.07 | 0.06 | 0.05 | 0.13 | 0.13 | 0.14 | 0.14 |
| | | 0.07 | 0.08 | 0.06 | 0.07 | 0.22 | 0.2 | 0.16 | 0.16 |
| | SCIHC011 | 0.06 | 0.06 | 0.05 | 0.05 | 0.15 | 0.15 | 0.13 | 0.13 |
| | | 0.06 | 0.06 | 0.06 | 0.05 | 0.16 | 0.15 | 0.14 | 0.14 |
| | SCIHC014 | 0.05 | 0.06 | 0.06 | 0.04 | 0.2 | 0.13 | 0.12 | 0.12 |
| | | 0.06 | 0.06 | 0.03 | 0.03 | 0.15 | 0.21 | 0.09 | 0.09 |

**Table 1** Percentages for SC-IO anterograde labeling per subnuclei.
